# A cell atlas of the adult *Drosophila* midgut

**DOI:** 10.1101/410423

**Authors:** Ruei-Jiun Hung, Yanhui Hu, Rory Kirchner, Fangge Li, Chiwei Xu, Aram Comjean, Sudhir Gopal Tattikota, Wei Roc Song, Shannan Ho Sui, Norbert Perrimon

## Abstract

Studies of the adult *Drosophila* midgut have provided a number of insights on cell type diversity, stem cell regeneration, tissue homeostasis and cell fate decision. Advances in single-cell RNA sequencing (scRNA-seq) provide opportunities to identify new cell types and molecular features. We used inDrop to characterize the transcriptome of midgut epithelial cells and identified 12 distinct clusters representing intestinal stem cells (ISCs), enteroblasts (EBs), enteroendocrine cells (EEs), enterocytes (ECs) from different regions, and cardia. This unbiased approach recovered 90% of the known ISCs/EBs markers, highlighting the high quality of the dataset. Gene set enrichment analysis in conjunction with electron micrographs revealed that ISCs are enriched in free ribosomes and possess mitochondria with fewer cristae. We demonstrate that a subset of EEs in the middle region of the midgut expresses the progenitor marker *esg* and that individual EEs are capable of expressing up to 4 different gut hormone peptides. We also show that the transcription factor *klumpfuss* (*klu*) is expressed in EBs and functions to suppress EE formation. Lastly, we provide a web-based resource for visualization of gene expression in single cells. Altogether, our study provides a comprehensive resource for addressing novel functions of genes in the midgut epithelium.

## Introduction

The adult *Drosophila* midgut, like its mammalian counterpart, is a complex tissue composed of various cell types performing diverse functions such as digestion, absorption of nutrients, and hormone production. Enterocytes (ECs) secrete digestive enzymes, absorb and transport nutrients, whereas enteroendocrine cells (EEs) secrete gut hormones that regulate gut mobility and function in response to external stimuli and bacteria. The midgut is a highly regenerative organ that has been used extensively in recent years as a model system to characterize the role of signaling pathways that coordinate stem cell proliferation and differentiation in the context of homeostasis and regeneration. For example, EGFR, JAK/STAT and Hippo signaling control intestinal stem cells (ISCs) growth and proliferation (Jiang et al., 2011; Jiang et al., 2009; Karpowicz et al., 2010; Shaw et al., 2010), while Notch signaling regulates ISCs differentiation (Micchelli and Perrimon, 2006; Ohlstein and Spradling, 2006, 2007). To maintain homeostasis, ISCs proliferate and give rise to a transient progenitor, the enteroblast (EB), defined by the expression of *Su(H)-lacZ*, a Notch pathway activity reporter (Micchelli and Perrimon, 2006; Ohlstein and Spradling, 2006). Both ISCs and EBs express the SNAIL family transcription factor, *escargot* (*esg)*. Most EBs differentiate into polyploid ECs, characterized by the expression of *Myosin31DF* (*Myo1A*) and *nubbin* (also called *pdm1*), whereas about 10% of EBs differentiate into EEs, marked by the expression of *prospero* (*pros*) (Figure 1A) (Micchelli and Perrimon, 2006; Ohlstein and Spradling, 2006). Recently, it was reported that new EEs in the adult are generated from ISCs through a distinct progenitor, called pre-EE, that expresses Piezo, a cation channel that senses mechanical tension (He et al., 2018; Zeng and Hou, 2015). In addition, the midgut is surrounded by visceral muscles, which control midgut movements and secrete many niche signals, such as Wingless (Wg), the EGFR ligand Vein (Vn) and the JAK-STAT ligand Unpaired1 (Upd1) to control ISC activities (Biteau and Jasper, 2011; Jiang et al., 2011; Lin et al., 2008).

**Figure 1.**
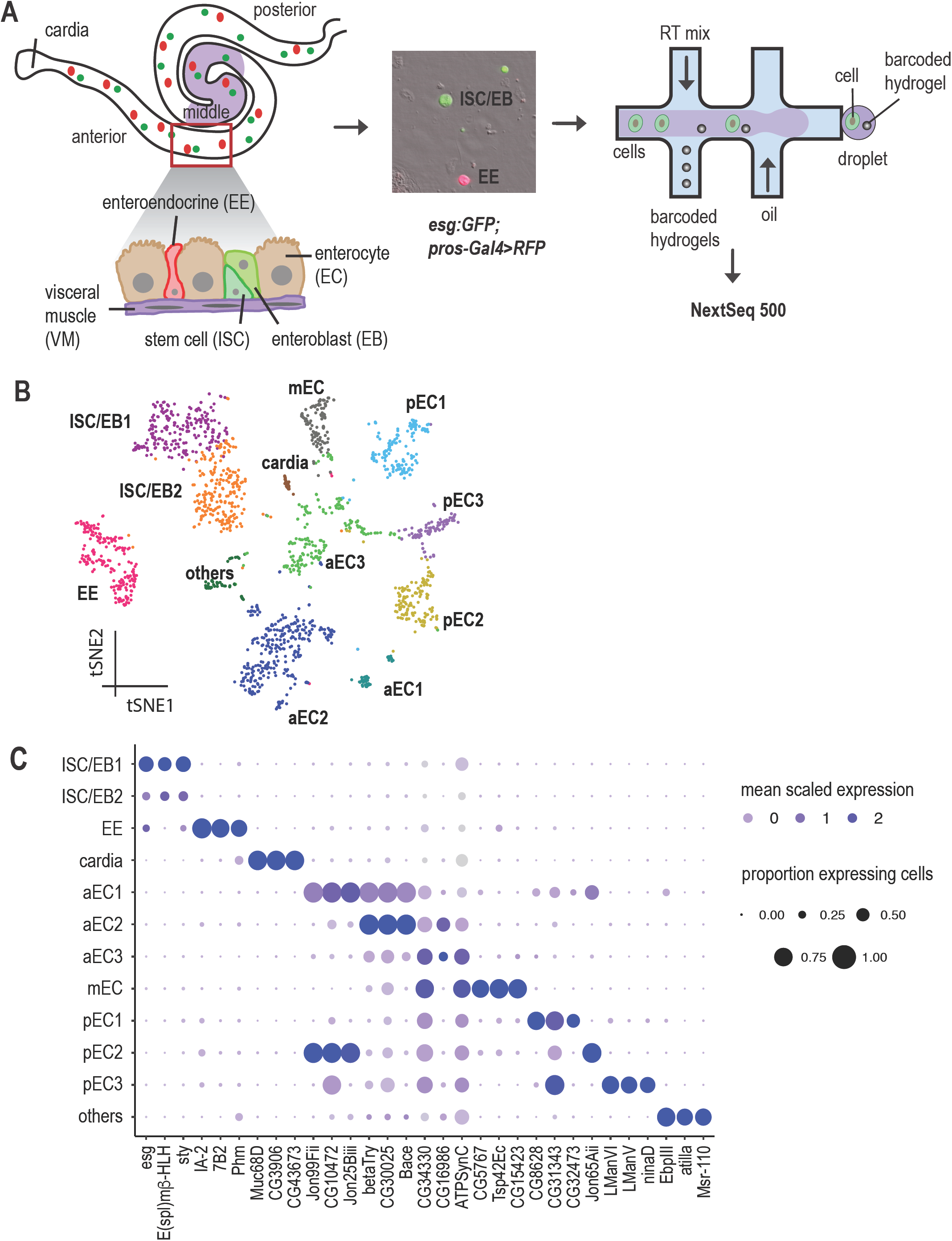
Single cell expression survey of the adult intestinal gut. A. Experimental design. Different regions of the midgut and different cell types are shown. For simplicity, we distinguish three (anterior, middle and posterior) regions of the midgut. Green, ISC/EB; red, EE. Cells were isolated and encapsulated using inDrop. B. Annotated cell types visualized on the t-SNE plot of 1753 cells. ISC/EB, intestinal stem cells/enteroblast; EE, enteroendocrine; aEC, anterior enterocyte; mEC, middle enterocyte; pEC, posterior enterocyte. C. Expression levels and the percentage of cells expressing the top 3 markers in each cluster is shown as a dot plot.

Like the compartmentalized mammalian digestive tract, the fly midgut can be divided into regions with distinct morphological, histological and genetic properties (Buchon et al., 2013; Marianes and Spradling, 2013). For example, the middle region of the midgut, which contains a group of specialized copper cells, is acidic and resembles the mammalian stomach (Dubreuil, 2004). In addition, EEs produce at least nine gut hormone peptides: Allatostatins (AstA, AstB/MIP, AstC), Tachykinin (Tk), neuropeptide F (NPF), DH31, CCHa1, CCHa2 and Orcokinin B (Chen et al., 2015; Reiher et al., 2011; Veenstra et al., 2008; Veenstra and Ida, 2014). These hormones are produced in specific regions. For example, AstA-producing EEs are located in the posterior region of the gut, whereas EEs in the anterior, middle and first half of the posterior midgut produce AstC (Veenstra et al., 2008). Moreover, individual EEs are able to produce combinations of different hormones. In particular, some NPF-producing EEs also produce Tk (Song et al., 2014; Veenstra et al., 2008). While differentiated cells, ECs and EEs function differently in different regions and show transcriptional differences, it is unclear whether these regional differences are also discernible in undifferentiated ISCs.

To further characterize gene expression and cell types in the adult fly midgut, we used single-cell RNA sequencing (scRNA-seq), as it provides an unbiased approach to survey cell type diversity and function, as well as to define relationships between cell types (Tusi et al., 2018; Wagner et al., 2018). Our study reveals novel molecular markers for each cell type; cell type specific organelle features; regional differences among ECs; a new subset of EEs; a transitional state from ISC to EC; and changes in gene expression during EC and EE differentiation. We demonstrate how the data set can be used to characterize new genes involved in gut cell lineages and in particular, we demonstrate that the transcription factor *klumpfuss* suppresses EEs formation. Finally, we built a web-based resource (https://www.flyrnai.org/scRNA/) that allows a user to query the expression of any genes of interest and compare the expression of any two genes in individual cells. Altogether, our study provides a valuable resource for future study of the *Drosophila* midgut.

## Results

### Unbiased single-cell transcriptomics identifies 12 distinct clusters in the adult *Drosophila* midgut

We used the inDrop method (Klein et al., 2015) to profile the transcriptome of 1753 midgut epithelial cells from 7-day-old females expressing GFP in progenitors, i.e. ISCs and EBs, and RFP in EEs (*esg:GFP/+; pros-Gal4,UAS-RFP/+*) (Figure 1A). Next, we used the Seurat algorithm to identify highly variable genes, performed linear dimensional reduction, determined statistically significant principal components, and generated unsupervised graph-based clustering (Satija et al., 2015). The inDrop run statistics are summarized in Table S1. The analysis revealed 12 distinct clusters that can be visualized using a t-distributed stochastic neighbor-embedding (tSNE) plot (Hinton, 2008) (Figure 1B). Each cluster was assigned to a specific cell type based on known specific markers (markers listed in Table S2, Figure S1). To facilitate mining of the data sets, we developed a visualization web portal (www.flyrnai.org/scRNA/), which allows users to search for either a particular gene or view a comparison of the expression of two genes across different clusters.

ISC/EB progenitors were assigned to two clusters based on the expression of *Dl* and *esg* (Figure S1B-C). Further analysis of these two clusters, based on detection of the downstream targets of Notch signaling, *Su(H)* and *Enhancer of split* complex *E(spl)-C*, did not allow us to distinguish ISCs from EBs (Bardin et al., 2010). Thus, we refer to these two clusters as ISC/EB1 and ISC/EB2. EEs were assigned to a single cluster based on the expression of *pros* (Figure S1D). Eight different clusters mapped to ECs from distinct gut regions based on the expression of different *Trypsin* genes (Buchon et al., 2013; Marianes and Spradling, 2013). We refer to the last cluster as “others”, as it might correspond to visceral muscle cells (VMs), trachea cells, enteric neurons and/or hemocytes, which are physically associated with the midgut (Figure S1G). Interestingly, three of the eight EC clusters, aEC1-3, could be mapped to the anterior region of the midgut because they express *alphaTry, betaTry, gammaTry, deltaTry,* and *epsilonTry* (Figure S2B-F). Another cluster, mEC, mapped to the middle region of the midgut based on the regional expression of *lab* (Figure S1E) and *ptx1* transcription factors (Dubreuil et al., 2001; Hoppler and Bienz, 1995; Marianes and Spradling, 2013; Strand and Micchelli, 2011), as well as *Vha100-4*, a component of Vacuolar H+ ATPase required for acid generation (Overend et al., 2016). Three additional clusters, pEC1-3, mapped to the posterior midgut based on *kappaTry, lambdaTry, iotaTry, etaTry,* and *zetaTry* expression (Figure S2H-L). Finally, the last EC cluster mapped to cardia secretory cells, which synthesize and secrete the peritrophic membrane that lines and protects the gut, based on the expression of *Pgant4* (Zhang et al., 2014) (Figure S1F). The expression of the top three markers for each cluster is shown in a dot plot (Figure1C). EEs represented 11% of the cells (compared to 10% expected (Beehler-Evans and Micchelli, 2015)). However, ISCs and EBs were over-represented (25% observed compared to 10% expected (Ohlstein and Spradling, 2006)), possibly due to the fact that ISCs/EBs have loose adherens junctions and are easier to dissociate from the epithelium (Baumann, 2001; Chiwei Xu, 2018). In addition, ECs were under-represented (60% compared to 80% expected), most likely due to our isolation method, which involved passing cells through two different sizes of cell strainers to minimize aggregates and doublets (ECs are much bigger than ISCs and are often lost when passing through a cell strainer).

To evaluate the robustness of the marker genes identified by the Seurat algorithm, we compared marker genes identified by Seurat to known markers reported in the literature based on antibody stainings, GFP reporters, *LacZ* reporters, Gal4 and enhancer trap lines, etc. (see full list in Table S3). Seurat was able to recover 74% of known markers in the major cell types (29 out of 39), including recovery of ∼80% of ISCs (17 out of 21) and EEs (11 out of 14) markers. To improve the recovery of marker genes, we added genes that met any one of the following three criteria: (1) The gene is expressed in more than 10 cells and is expressed in >60% of cells in one cluster but less than 20% of cells in other clusters; (2) The gene is expressed in more than 10 cells and the gene is expressed in >80% of cells in multiple clusters and >20% of cells in all clusters and (3) The gene is expressed in a small number of cells (3-10 cells) and all of the cells that express the gene are in the same cluster. These criteria were chosen based on testing parameters to recover additional known markers. Combining this approach with Seurat, we were able to recover 85% of well-known markers (33 out of 39) in the literature, including 90% of ISC/EB (19 out of 21) and 86% of EE (12 out of 14) markers (Table S3), representing a very low false negative rate. Finally, we compared our data with cell-type preferentially expressed genes from the Flygut-seq resource (http://flygutseq.buchonlab.com/), a RNAseq study that separated cell types using fluorescence-activated cell sorting based on cell-type specific Gal4 activities (*Dl-Gal4* for ISCs, *Su(H)-Gal4* for EBs, *Myo1A-Gal4* for ECs, *pros-Gal4* for EEs and *24B-Gal4* for VMs). We defined cell type specific genes using Seurat algorithm, based on the highly variable expression in different cell types. We found that 89% of ISCs/EBs (p-value, 2.1E-14), 90% of EEs (p-value, 4.4E-15), and 61% of ECs (p-value, 5.4E-15) specific genes from our scRNA-seq data set overlap with the cell-type preferentially expressed genes in Flygut-seq (Figure S3A).

### Gene expression signatures of each cluster

Next, we analyzed the gene expression signatures of each cluster and determined the enrichment of specific groups of genes involved in distinct cellular processes using the Gene List Annotation for *Drosophila* (GLAD) online resource (Hu et al., 2015). Genes categorized as cell cycle, cytoskeletal proteins, major signaling pathways, RNA-binding, spliceosome and transcription factors are enriched in ISC/EB progenitor cells, whereas genes involved in metabolic process, serine proteases and transporters are enriched in ECs (Figure 2A, p-value in Table S4). Detailed analyses of the expression of canonical genes of signaling pathways revealed that components of the EGFR, Hippo, Imd, insulin and Notch pathways are enriched in progenitors cells (especially in the ISC/EB1 cluster), and that the Imd and Toll immune pathways are enriched in a subset of ECs, pEC2 cluster (Figure 2B). Further analyses showed that different EC clusters are enriched for different types of cellular processes. For example, cardia ECs are enriched for genes involved in glycan biosynthesis, consistent with the known role of cardia cells in secretion of glycosylated proteins (Zhang et al., 2014), and posterior ECs are enriched for genes involved in lipid metabolism (Figure 2C) (Marianes and Spradling, 2013). In addition, genes encoding endoplasmic reticulum and Golgi stack components are enriched in cardia ECs, consistent with their roles in polarized secretion of peritrophic matrix and mucin (Golgi staining in cardia cells is shown in (Zhang et al., 2014)). Interestingly, ribosomal proteins are enriched within ISC/EB1 and 2 (Figure 2D, 3A-B’’ and 3D) consistent with the high abundance of ribosomes in ISCs giving them a darker appearance in electron micrographs compared to ECs (Miller, 1950; Ohlstein and Spradling, 2006) (Figure 3A-A’’). The majority of ribosomes in ISCs are free ribosomes. ER-bound ribosomes require signal recognition particle (SRP) binding to facilitate their docking onto the ER through SRP receptors and components of SRPs are expressed at low levels in ISCs and enriched in differentiated ECs (20% of ISCs expressed SRPs vs 36% of ECs, Figure 3E and Table S5), further supporting our observation that ISCs contain more free ribosomes (Figure 3A-B’’). Interestingly, reduction in ribosome levels impairs the lineage commitment in human hematopoiesis (Khajuria et al., 2018). Another characteristic of stem cells is that they often use glycolysis rather than oxidative phosphorylation as their energy source due to immature mitochondria (fewer cristae) (Figure 3B’-B’’) [(Teixeira et al., 2015) and reviews in (Hu et al., 2016)]. Consistent with this, we observe fewer mitochondrial cristae in ISCs than that in ECs (Figure 3B’-B’’). During differentiation, ATP synthase (complex V) promotes the maturation of mitochondrial cristae through dimerization and up-regulation of the ATP synthase complex (Teixeira et al., 2015). We observe an enrichment of ATP synthase complex V in cluster aEC3, mEC, and pEC1 (Figure 2D). In particular, cluster aEC3 also retains some characteristics of ISCs that are attributable to ribosome enrichment; therefore, we speculate that aEC3 are relatively premature ECs and are in the process of differentiation (see also results from the reconstruction of lineage differentiation trajectories analysis below). Finally, proteins enriched in synaptic vesicles are observed in EEs and proteins enriched in brush border are observed in ECs (Figure 2D and 3C-C’).

**Figure 2.**
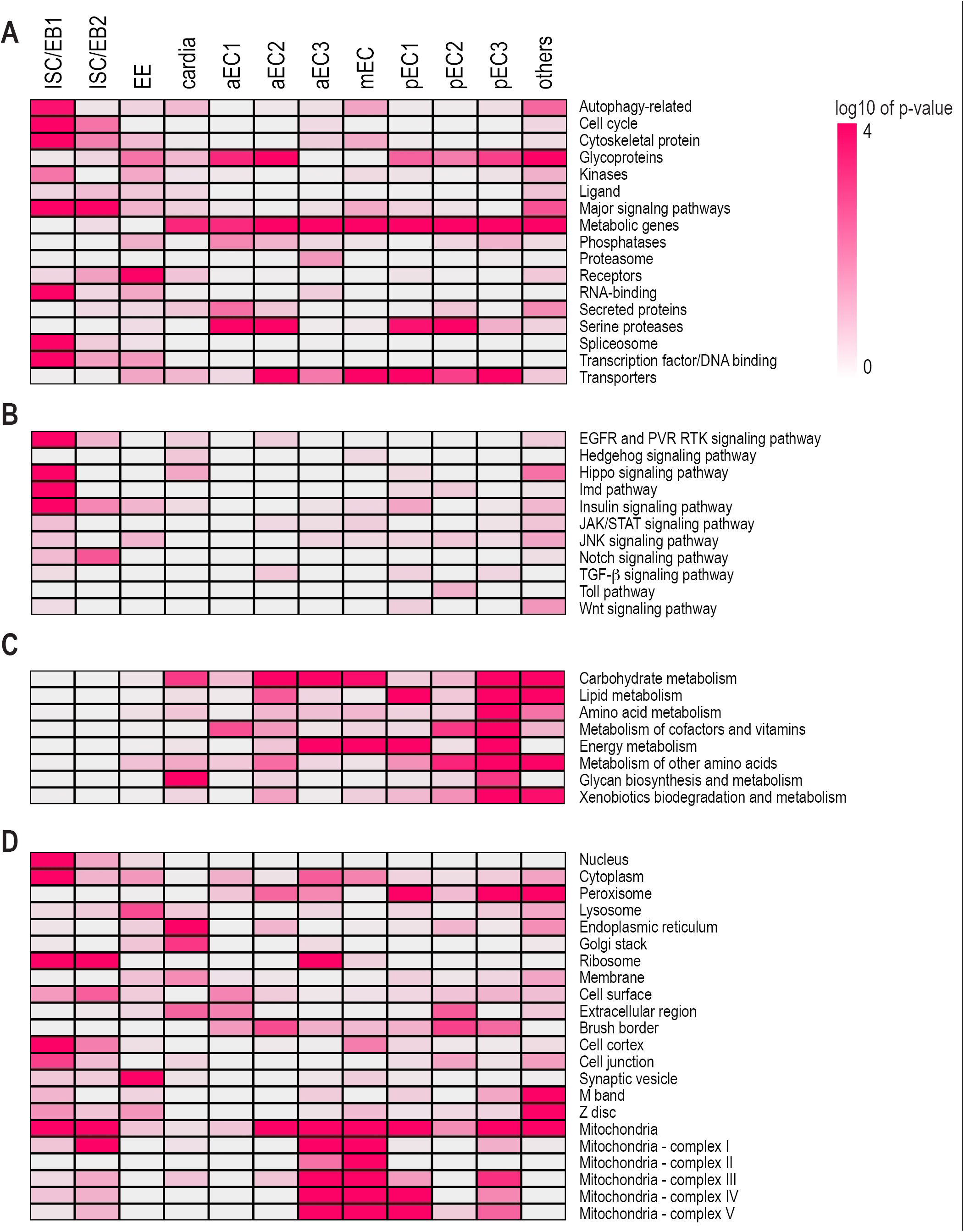
Gene set enrichment analysis. Markers identified by Seurat and additional criteria in different clusters are categorized using GLAD (A); transcriptional target genes of major signaling pathways (B); metabolic pathways (C); cellular compartments (D).

**Figure 3.**
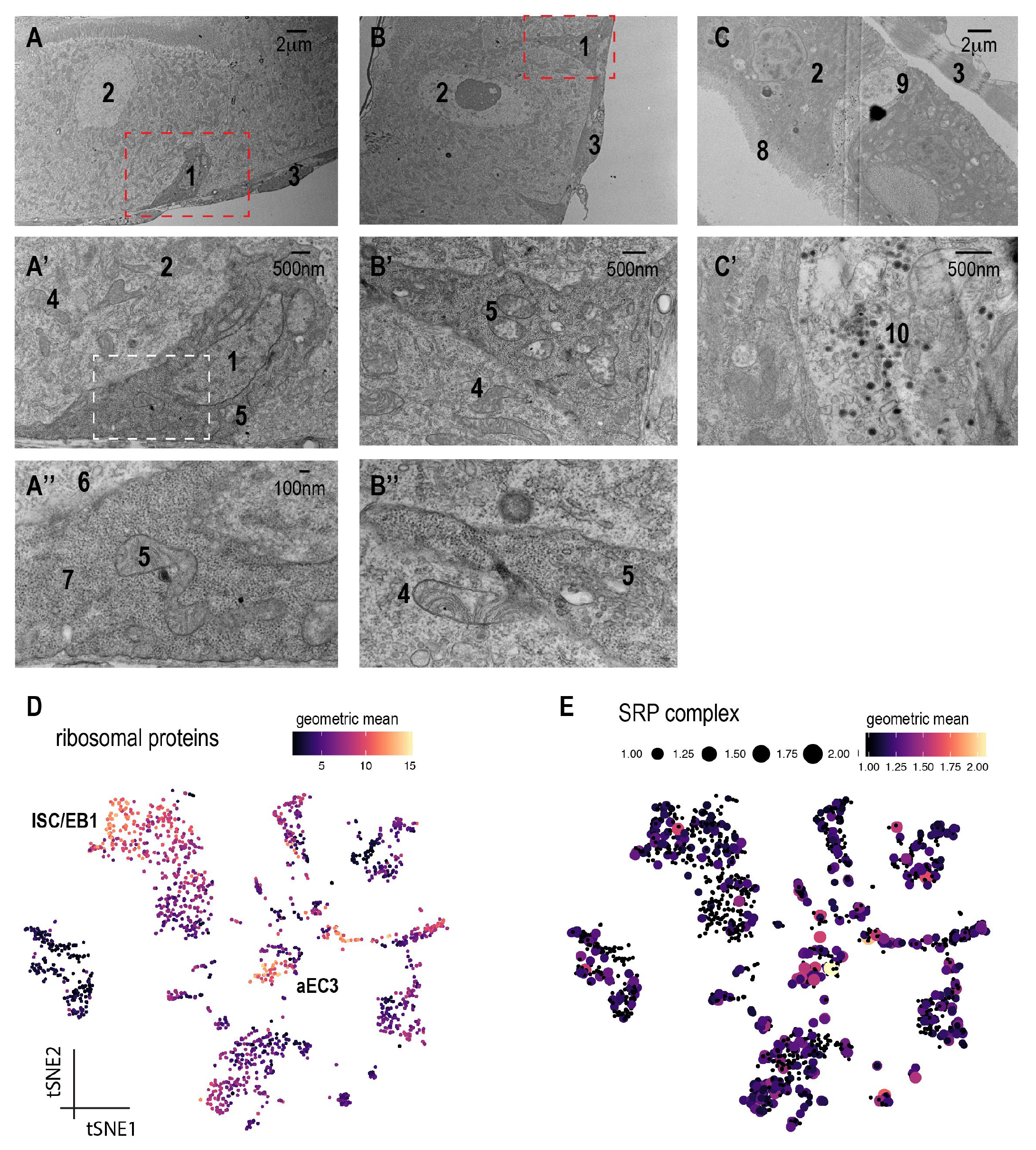
Electron micrographs of the ultrastructure of different cell types. A. ISC (1) with darker density than its neighboring EC (2) resides on the basal side of the epithelium. They are often triangular shape with extensive basement membrane contact. A’. Magnified view of A outlined by the red dash box. A’’. Magnified view of A’ outline by the white dash box. The density of ribosomes (7) in ISCs is more pronounced than that of ribosomes (6) compared to ECs. B’ is a magnified view of B. Mitochondria in ISCs (5) contain less cristae than mitochondria of ECs (4), which can be visualized better in B’’. Mitochondria in ISCs look “empty”. C. EEs (9) contain many “loaded” secreted vesicles (10, dark vesicle), which can be better visualized in C’. 1. ISC; 2. EC; 3. visceral muscle; 4. mitochondria in EC; 5. mitochondria in ISC; 6. ribosomes in EC; 7. ribosomes in ISC; 8. microvilli; 9. EE; 10. secretory vesicles. D. tSNE plot of the geometric mean of ribosomal proteins. Ribosomal genes were expressed at high level in the ISC/EB1 and aEC3 cluster. E. tSNE plot of the geometric mean of the SRP complex, a main component for ER-bound ribosomes. The SRP complex (consisting of 6 proteins) is expressed at low level in ISCs/EBs but at high level in ECs, indicating that the majority of ribosomes in ISCs/EBs are free ribosomes and not ER-bound ribosomes.

### Regional variation along the gut

Cell morphology and digestive functions are different along the length of the *Drosophila* midgut (Marianes and Spradling, 2013). For example, cells in the middle midgut secrete acid and absorb metal ions, whereas cells in the posterior midgut are enriched with lipid droplets and absorb lipid nutrients. These characteristics reflect regionalized gene expression differences as shown by bulk RNA-seq studies (Buchon et al., 2013; Marianes and Spradling, 2013). We searched for differentially expressed transcription factors that could underlie regionalized gene expression and identified a number of potential candidates. The genes *vnd* and *odd* are preferentially expressed in the anterior region. Interestingly, *vnd* and *odd* are expressed in the anterior and posterior midgut in the embryo, but the regionalized expression pattern in the adult midgut needs future investigation (Jimenez et al., 1995; Ward and Coulter, 2000). The genes *lab, Ptx1, CREG, apt* and *dve* are preferentially expressed in the middle midgut. The homeobox genes *lab, Ptx* and *dve* have been shown to express in the adult middle midgut (Buchon et al., 2013; Dutta et al., 2015; Strand and Micchelli, 2011). Lastly, the genes *bab2, ham, cad, Ets21C, Hnf4* and *hth* are preferentially expressed in the posterior midgut. The homeobox gene *cad* and *Ets21C* have been reported to be expressed in the posterior midgut (Buchon et al., 2013; Choi et al., 2008; Jin et al., 2015). *Hnf4* is expressed in the midgut in the developing embryo; whether it is expressed in the adult posterior gut is not known (Zhong et al., 1993). The *hth* gene encodes a cofactor of Ubx, which is known to regulate *Dpp* in the middle midgut; the expression pattern of *hth* is not known (Li et al., 2016). The regionalized expression and expression patterns of these candidates from flyatlas are summarized in Table S6.

Stem cell morphology and proliferation activity also differs in different regions of the gut. Although previous cell-specific RNAseq revealed regional differences in stem cell transcriptomes, we were not able to identify subgroups or regional ISC/EB clusters (Dutta et al., 2015), despite the fact that that some stem cells expressed some regional markers, such as *lab* or *Ptx1*. In addition, based on the fact that *Dl* is expressed at higher levels in the anterior region than posterior region (Dutta et al., 2015), it is possible that ISC/EB1 is from the anterior region and ISC/EB2 from the posterior region. Alternatively, It is possible that the regional differences in ISCs transcriptomes are less prominent than the regional differences in ECs transcriptomes since EC clusters are separated by regions (Buchon et al., 2013; Marianes and Spradling, 2013).

### Reconstructing lineage differentiation trajectories

To investigate the relationship between different clusters and explore the process of cell differentiation, we used single-cell trajectories reconstruction, expression and mapping (STREAM) to arrange cells in a pseudotime trajectory (Huidong Chen, 2018). We specified ISC/EB as the start state and STREAM arranged cells as a function of pseudotime, resulting in three branches (Figure 4A). The ISC/EB1 cluster in this analysis is more closely related to the ISC state because of its position in pseudotime earlier than ISC/EB2 (Figure 4A, subway plot). Following ISCs/EBs, cells are arranged into three branches: one corresponding to EEs; another corresponding to ECs; and a third branch, barely connected to ISCs/EBs, corresponding to other cell types (Figure 4B, stream plot; this is expected because those cells are not derived from ISCs). Based on their organelle features of ribosomes enrichment, we predicted that aEC3 are premature ECs that have undergone the transition from ISCs/EBs to ECs. Indeed, the aEC3 cell population in the subway and STREAM plot is located at the branch between ISCs/EBs and ECs, indicating that aEC3 is relatively premature than other EC clusters (Figure 4A-B, light green). Although ECs from different regions of the gut reside in a single branch along different pseudotime, we cannot infer specific lineage relationships among ECs from different regions. For example, the observation that aECs and mECs are at an earlier pseudotime than pECs does not imply that aECs give rise to pECs. It is possible that pECs expressed more various digestive enzymes such as trypsins, maltases, Jonah proteases and mannosidase, so STREAM thought pECs are generally more mature than aECs and mECs. In addition, *Dl*, an ISC marker gene, is expressed at high levels at the starting point (S2) and decreases in expression level over time in differentiating EEs and ECs (Figure 4C), indicating that STREAM has built accurate trajectories. Interestingly, as in the case of the Seurat analysis, we still cannot resolve a clear separation between ISCs and EBs. We also discovered genes with patterns similar to *Dl*, such as *MRE16* (lncRNA:CR34335) and *klumpfuss* (*klu*) (Figure 4D and G). STREAM also discovers diverging genes, ie., genes expressed differentially between bifurcating branches. For example, *α-trypsin* and *7B2* are differentially expressed between EE and EC bifurcating branches (Figure 4E and F, S1-S3 vs. S1-S4). We also identified *7B2* as an EE marker; its mammalian ortholog is a secreted chaperon protein expressed in neuroendocrine cells (Helwig et al., 2013). In addition, the expression of septate junction protein complex, *mesh, ssk* and *Tsp2A,* gradually increased as ISCs differentiate to ECs or EEs, consistent with the observation that septate junction complex proteins are present in the apicolateral regions between ECs and EEs, but not ISCs (*mesh*, Figure 4I) (Chiwei Xu, 2018; Izumi et al., 2016; Izumi et al., 2012; Yanagihashi et al., 2012).

**Figure 4.**
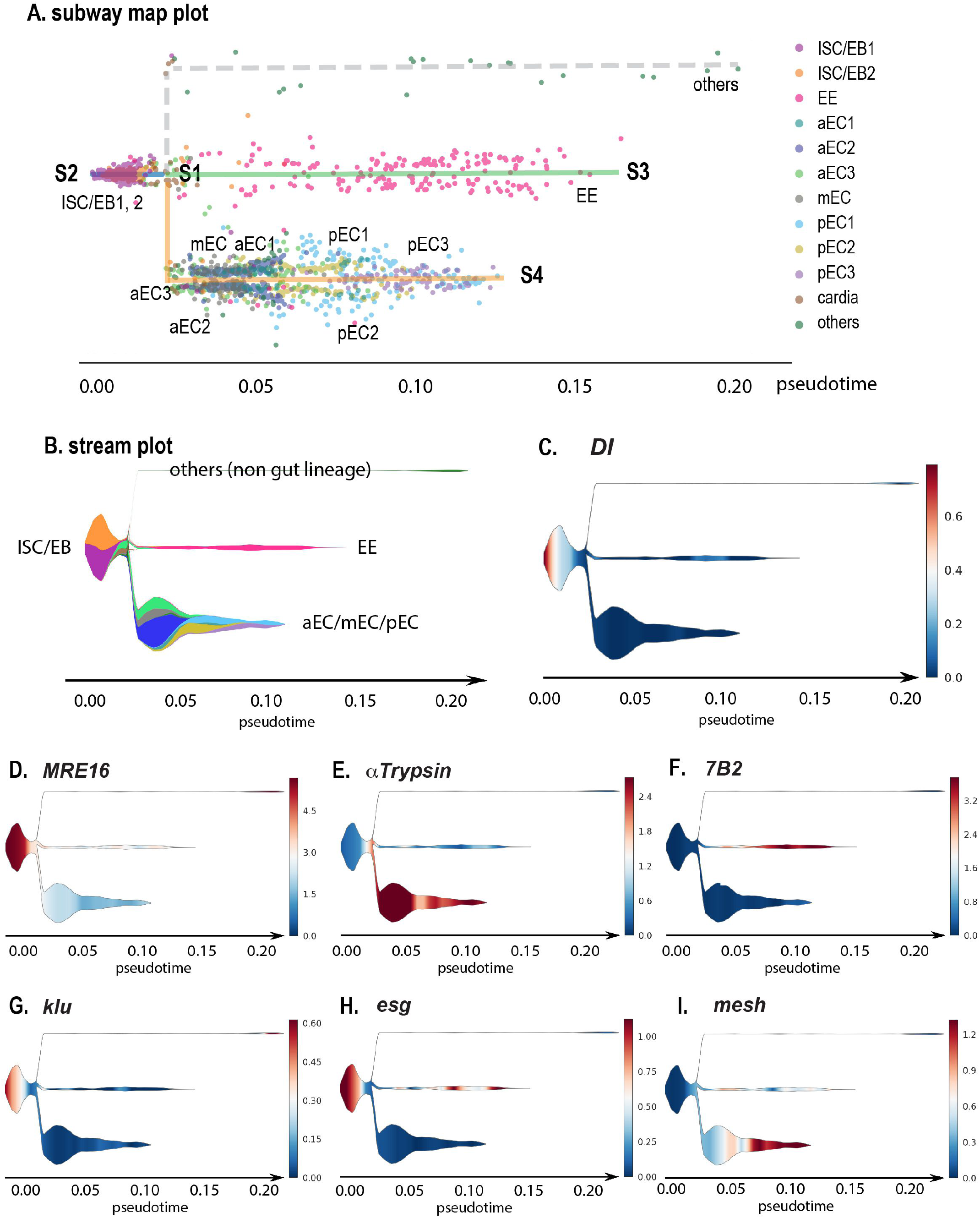
Single-cell trajectories reconstruction, exploration and mapping (STREAM) analysis. A. Subway map plot. Each cell type is color-labeled. S1-S4 indicates cell states. S2 was selected as the initial state. B. STREAM plot: cell types and cell densities are shown along different trajectories at a given pseudotime. The width of each branch is proportional to the total number of cells. Cell type color is the same as in A. C. Stream plot of *Dl* gene expression. The expression of *Dl* is restricted to the initial state. D. STREAM plot of *MRE* 16, a non-coding RNA. Its expression pattern is restricted to ISC/EB and similar to *Dl*. E. STREAM plot of *αTrypsin*, a diverging gene as its expression is different between S1-S4 (EC) and S1-S3 (EE) branches. F. STREAM plot of *7B2*, also a diverging gene between S1-S3 (EE) and S1-S4 (EC). G. STREAM plot of *klu*, a new marker identified with similar expression pattern to *Dl*. H. STREAM plot of *esg*. I. STREAM plot of *mesh*.

### Identifying new progenitor markers

Transcription factors play important roles in regulating stem cell maintenance and differentiation. Our scRNA-seq analysis identified many previously known transcription factors (*esg, E(spl)mβ-HLH, myc, bun, E(spl)m3-HLH, N, Peb, fkh, sox21a, cic, Rel, HmgD, msn*), polarity and adhesion proteins (*mira, mew, shg, mys, LanA, Nrg*) and other molecules (*robo2, hdc, Ets21C*, etc.) important for stem cell proliferation and maintenance (see Table S7 for top 20 transcription factors and Table S8 for the full list). We also newly identified additional transcription factors that are enriched in the ISCs/EBs cluster: *Df31, Blimp-1, mamo, Sox100B, klu, lola, Eip75B, nej, elB, Smr,* and *fs(1)h*. Interestingly, although the expression of *klu* as a function of pseudotime is similar to *Dl* in the STREAM analysis, we found an opposite (mutually exclusive) expression pattern of *Dl* and *klu* in individual cells using our web-based resource. We confirmed that *Klumpfuss* (*klu*) is expressed in the EBs, as its expression co-localized well with the EB reporter *Su(H)-LacZ* (Figure 5A-C). *Klu* is a Zinc finger transcription factor, previously characterized as a regulator of self-renewal in *Drosophila* neuroblasts (Berger et al., 2012; Xiao et al., 2012). During hematopoiesis, *Klu* acts downstream of Notch and Lozenge (Lz) to promote crystal differentiation (Terriente-Felix et al., 2013). Knockdown *of klu* with two independent RNAi lines in ISCs and EBs with *esgGal4-tubgal80^ts^* in the adults for 7 days resulted in an increase of the EE population (Figure 5D-G), which is similar to the *Notch* loss of function phenotype (Micchelli and Perrimon, 2006; Ohlstein and Spradling, 2006). We observed an increase in the number of AstA expressing EEs when knocking down *klu* in progenitor cells (Figure 5H), consistent with an increase of AstA EEs in *Notch* loss of function MARCM clones (Beehler-Evans and Micchelli, 2015). We also confirmed that *lola*, a transcription repressor antagonizing Notch (Zheng and Carthew, 2008), is expressed in *esg+* cells, but only in the middle region of the midgut (Figure S4A).

**Figure 5.**
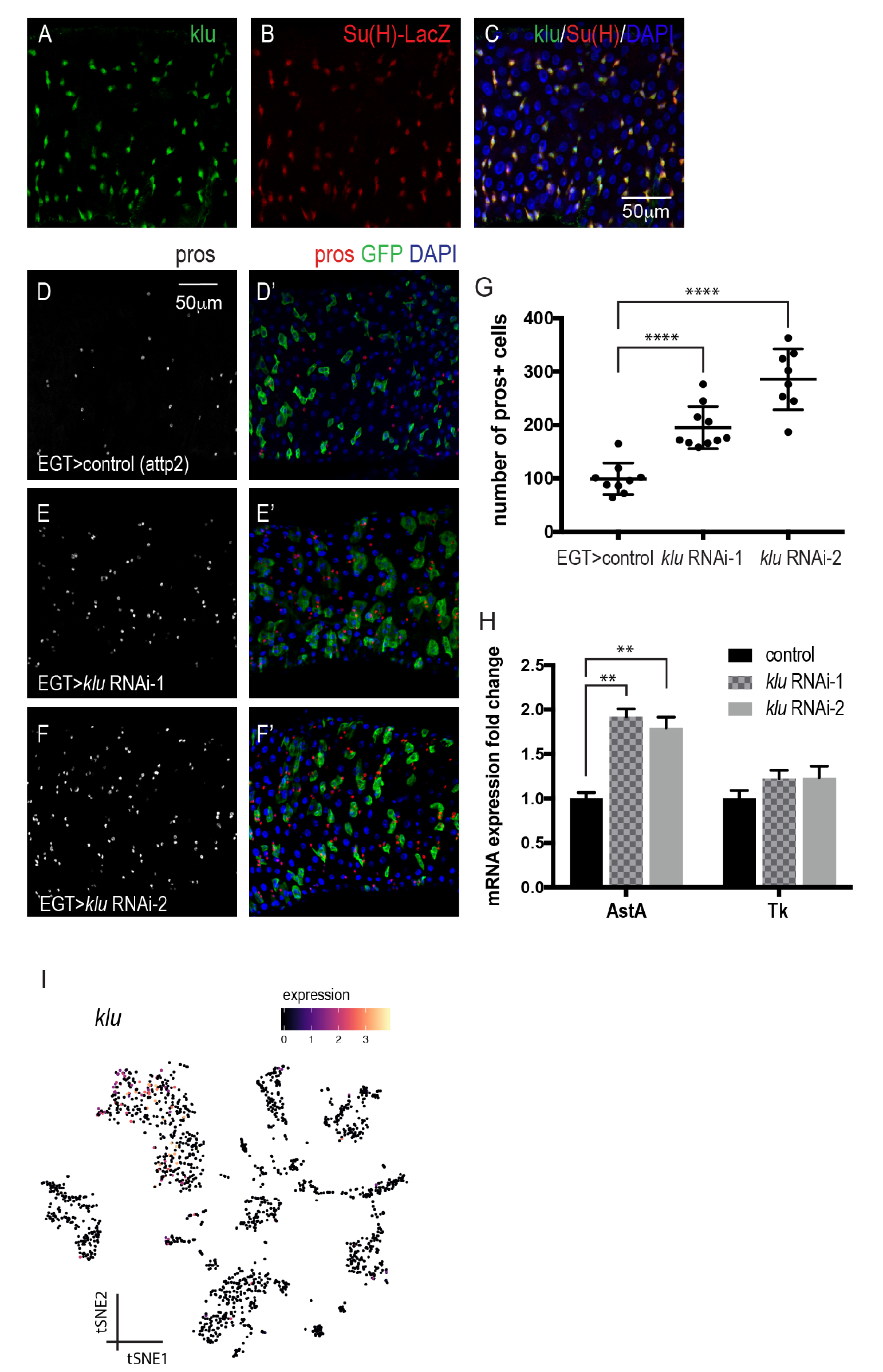
*klu* expression in EBs and its loss of function phenotype. A-C. Co-expression of *klu* and the EB reporter *Su(H)-LacZ*. D-F’. Knockdown of *klu* in ISCs/EBs results in an increase of EEs, marked by *pros* staining. red, *pros*; green, *esg*; blue, DAPI. G. Quantification of EEs number. Data are represented as mean ± SEM. Two tailed t-test, ****, p<0.0001. H. RT-qPCR measurement of *AstA* and *Tk* expression from midguts expressing *klu* RNAi in adult ISCs/EBs for 7 days. *rp49* is used for normalization. Data are represented as mean ± SEM. Two tailed t-test, p=0.0011 for control vs. RNAi-1; p=0.0047 for control vs. RNAi-2. I. tSNE plot of *klu* expression level across different clusters.

In addition to transcription factors, we also confirmed that *leucine-rich-repeats and calponin homology domain protein* (*Lrch*), *Zipper* (*zip*) and *inscutable* (*insc)* are expressed in *esg+* progenitor cells (Figure S4B-D) in different regions of the midgut. Interestingly, all of them are related to the cytoskeleton, suggesting that the cytoskeleton plays important roles in stem cell shape and biology, consistent with the results of GLAD enrichment analyses (Figure 2A). The gene *Lrch*, which is expressed in both anterior and posterior regions of the midgut, encodes a scaffold protein well conserved across animals that interacts with actin to stabilize the cell cortex and position of the mitotic spindle during cell division (Foussard et al., 2010). *Zip* encodes cytoplasmic myosin II, which also binds to the actin cytoskeleton (Franke et al., 2006), and is co-expressed with *esg+* in cells in the middle of the midgut. *Insc*, which encodes a cytoskeletal adaptor protein required for apical-basal spindle orientation for asymmetric cell division in neuroblasts (Kraut et al., 1996), is co-expressed with *esg+* throughout the midgut, suggesting that it may be important for cell fate decisions.

### Identifying new EE markers and characterization of a subset of EEs

EEs are chemosensory cells that secrete regulatory hormones in response to luminal contents, such as nutrients and bacterial metabolites, to regulate gut physiology, food intake and glucose homeostasis. We detected all nine gut hormones in the EE cluster, *AstA, Mip, AstC, Tk, NPF, Dh31, CCHa1, CCHa2* and *Orcokinin*, as well as several key enzymes important for hormone biosynthesis, processing and secretion, such as *amon, Phm, Pal2, svr*, which are also targets of the transcription factor *dimmed* (Beebe et al., 2015; Hadzic et al., 2015; Reiher et al., 2011). We also identified markers enriched in the EE population, *IA-2, 7B2, nrv3, CG30183* and *unc-13-4A* and genes involved in vesicle docking and secretion, such as *Sytalpha, Sytbeta, Syt1, cac, nSyb* and *Syx1A* (Table S8).

We also noticed that *esg*, a marker for ISCs/EBs, is not only expressed in ISCs and EBs but also in a subset of EEs (*pros+*) (Figure S1). To validate this observation, we examined the expression of a GFP insertion in the *esg* locus and *pros-Gal4* driven RFP. In most cases, consistent with *esg* being a progenitor marker and *pros* a differentiation EE marker, *esg* and *pros* do not co-localize in individual cells. This mutually exclusive expression pattern is observed in the cells localized to the anterior and posterior region of the midgut. However, *esg+ pros+* double positive cells are detectable in the middle region and a small portion of the posterior midgut (Figure 6A-B). To further characterize these *esg+pros+* cells, we examined 196 EEs and the unsupervised clustering results in two sub-clusters: *esg+pros+* and *esg-pros+* cells. Unexpectedly, *esg+pros+* cells also express Tk and NPF hormones, suggesting that *esg+pros+* cells are mature EEs rather than EE progenitors (Figure S5B-D). We examined the expression of esg and NPF and found that they indeed colocalize in cells in the middle region of the midgut, but not in the anterior region (Figure 6C-D’’). Altogether, these analyses identify a new subset of EEs with different molecular properties.

**Figure 6.**
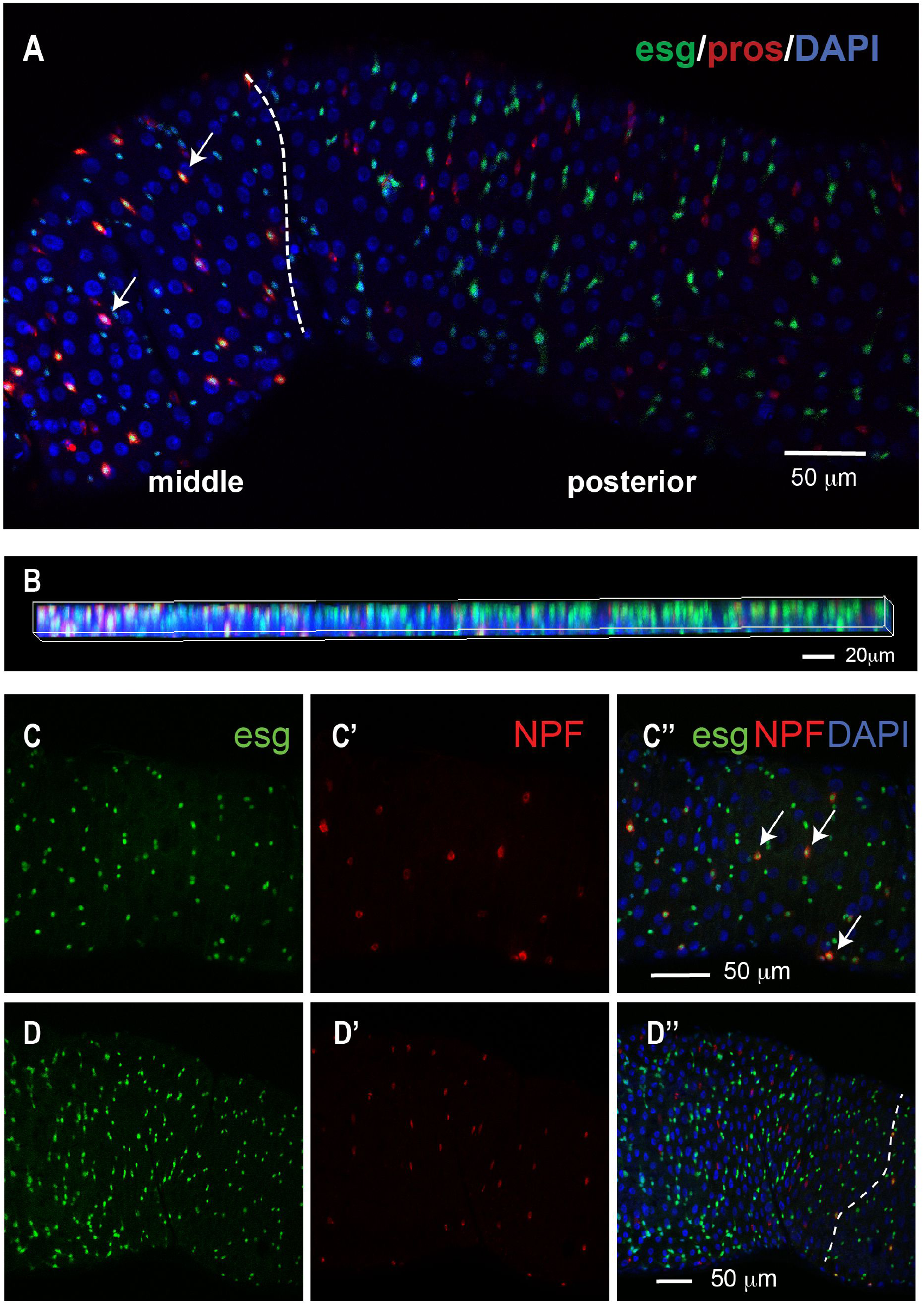
Characterization of a subset of EEs. A-B. The majority of *pros+ esg+* double positive cells are located in the middle of the midgut, while very few double positive cells are located in the posterior region. *pros*, a EE marker; *esg*, a progenitor marker. Dash line indicates the boundary between middle and posterior midgut regions. Arrow, *pros+esg+* double positive cells. B. Z stacking 2D image showing the colocalization of *pros* and *esg* in the middle but not the posterior region. Genotype, *esg-sfGFP/+, pros-Gal4; UAS-mCD8.RFP/+*. C-D. Image of middle (C-C’’) and anterior (D-D’’) midguts. Arrow, *NPF+esg+* double positive cells. Dash line indicates the boundary between anterior and middle midgut regions. Genotype: *esg-sfGFP/+, UAS-mcherryCAAX; NPF-Gal4/+*.

Individual EEs are known to express a combination of 2-3 gut hormones (Veenstra et al., 2008; Veenstra and Ida, 2014). For example, some EEs in the posterior midgut produce both AstA and AstC (Beehler-Evans and Micchelli, 2015). Interestingly, our scRNA-seq data reveals that AstA EEs are also able to produce CCHa1 and CCHa2, suggesting that individual EEs are capable of generating up to 4 different gut hormones (Figure S5A). This is consistent with the observation that CCHa1 EEs produce AstA and CCHa1 EEs are able to produce CCHa2 (Veenstra and Ida, 2014). In on our scRNA-seq data, 17 out of 46 AstA cells (37%) co-expressed AstC, CCHa1 and CCHa2. Thus, it is likely that individual EEs are able to express up to 4 different gut peptide hormones.

### ISCs and EBs share very similar transcriptomes

As our scRNA-seq analysis did not separate ISCs and EBs into two different clusters, we attempted to identify, among the ISC/EB1 and 2 clusters, cells with high-confidence ISC and EB markers, and then compared the difference between the two populations. For ISCs we selected cells expressing *Dl* and none of the Notch downstream targets, *E(spl)m2-BFM, E(spl)m3-HLH, E(spl)m4-BFM, E(spl)m5-HLH, E(spl)m7-HLH, E(spl)malpha-BFM, E(spl)mbeta-HLH, E(spl)mgamma-HLH* and *klu* (Housden et al., 2013). For EBs, we selected cells with no *Dl* expression and at least 3 of the 9 Notch downstream targets expression. This analysis allowed us to identify 26 ISCs and 73 EBs that we used to define ISC and EB transcriptomes. Next, we applied DESeq2 and additional criteria (mean expression >0.5 and log2 fold change >10) to identify genes specific to either ISCs or EBs. This analysis recovered *Dl* as a ISC marker, *E(spl)-C* and *klu* as EBs markers. While we did not observe additional genes specifically expressed at high levels in ISCs, a number of genes were preferentially expressed at high level in EBs, including *Rel, Dtg, l(2)gl* and some protease genes which are highly expressed in ECs (Table S9), suggesting that these EBs are already primed to express EC genes. Note that when we used another set of markers to define ISCs and EBs: *Dl+klu*-for ISC and *Dl-klu+* for EB as we identified *klu* as a new EB marker, similar results were obtained. Finally, principle component analysis (PCA) of ISCs and EBs showed that these two groups almost overlap completely (Figure S6), indicating that there is minimal difference between presumptive ISCs and EBs, which is also consistent with the PCA in Fig.1B from (Dutta et al., 2015).

### Comparison of fly midgut cell types to mammalian intestinal and airway tracheal epithelia

Recent scRNA-seq studies have provided a cell atlas of both the mammalian small intestine and tracheal epithelia (Haber et al., 2017; Montoro et al., 2018; Plasschaert et al., 2018). Cell types in these tissues are similar because they both contain stem cells, transit amplifying cells (TAs in intestine, club cells in trachea), endocrine cells, tuft and goblet cells. Based on the marker genes expression identified in the mammalian intestine, *Drosophila* ISCs are reminiscent of intestinal ISCs, TAs and immature distal ECs (Figure S3B) whereas *Drosophila* EEs are similar to EEs and tuft cells in the intestine. *Drosophila* cardia cells are similar to goblet cells, consistent with their role in secreting mucins. Mammalian Paneth cells are similar to *Drosophila* aEC2, possibly due to the expression of lysozymes in both cell types. Overall, *Drosophila* ECs are similar to mammalian intestinal ECs, however, the regional differences of *Drosophila* ECs do not match to the regional differences in the mammalian intestine (Figure S3B). P-values of these comparisons are summarized in Table S10.

In contrast to the mammalian intestine organized in crypts and villi, the *Drosophila* gut is composed of a single layer of cells (pseudostratified epithelium), which is more similar to the mammalian tracheal epithelium. Although the primary function is different, the role of stem cells to replenish all cell types is the same. Thus, we compared the similarity of stem cells and EEs, but not ECs and ciliated cells. *Drosophila* ISCs are reminiscent of tracheal basal stem cells (Figure S3C), whereas *Drosophila* EEs resemble basal stem cells, neuroendocrine cells, and ionocytes (a rare cell type that express the cystic fibrosis transmembrane conductance regulator (CFTR)) in the tracheal epithelium.

## Discussion

The *Drosophila* intestinal epithelium has been used to study stem cell regeneration and maintenance, as well as how signaling events from different cell types influence stem cell behaviors. Here, we surveyed the cell types of the adult intestinal epithelium using scRNA-seq and identified all known cell types, as well as one novel cell type (*esg*+*pros*+) in the middle region of the midgut. Our study recovered 90% and 86% of previously known ISCs and EEs markers, respectively, demonstrating the robustness of the scRNA-seq approach. Interestingly, gene expression analysis revealed that ISCs are enriched for free ribosomes and possess mitochondria with fewer cristae. We also identified new transcription factors and cytoskeletal proteins preferentially expressed in the ISCs/EBs population. In particular, we showed that *klu* is specifically expressed in EBs and colocalizes with the EB reporter *Su(H)-LacZ*, and showed that knockdown of *klu* in ISCs/EBs (with *esg-Gal4*) results in an increase of EEs, suggesting that *klu* inhibits EE differentiation.

In our scRNA-seq analysis, although we observed two ISC/EB clusters, we could not dichotomously distinguish ISCs and EBs. We observed that Notch downstream targets, *E(spl)m3-HLH, E(spl)malpha-BFM, E(spl)mbeta-HLH, E(spl)mgamma-HLH* and the new EB marker, *klu*, are expressed in both the ISC/EB1 and 2 clusters. In addition, we did not observe previously described regional differences in stem cell transcriptomes (Dutta et al., 2015; Marianes and Spradling, 2013). As we observe regional differences in ECs, it is possible that regional differences in ISCs are less prominent than in ECs. The other possibility is that regional difference genes in ISCs are expressed at low levels, which are not easily detected by scRNA-seq. To further attempt to distinguish ISCs and EBs we used STREAM; however, this analysis was also not able to separate these two cell states. To overcome these limitations, we defined ISC and EB transcriptomes by manually selecting cells based on high-confidence ISC and EB markers. PCA showed that the transcriptomes of ISCs and EBs are very similar. Differential gene expression analysis between these two cell populations allowed us to identify a small number of genes expressed higher in EB than ISC. Most of them are also expressed in ECs, suggesting that EB start to turn on EC differentiation genes.

Regarding EEs, we identified candidate markers and were able to identify a subset of EEs expressing NPF and Tk in the middle of the midgut that express the *esg* progenitor marker. In addition, we found that individual EEs are able to express up to four different hormones, in contrast to the view that these cells only produce two hormones (Song et al., 2014; Veenstra et al., 2008). Interestingly, a recent mammalian study showed that EEs modulate their hormonal repertoire in response to local cues (Beumer et al., 2018). For ECs, we were able to identify seven cell subpopulations, residing in the anterior, middle, and posterior of the midgut that express different types of trypsins, proteases and transporters. We also identified a subpopulation of ECs, aEC3, a relatively premature EC, supported by STREAM analysis and organelle characteristics.

Our study provides a resource to further characterize the molecular signature of each cell type and gene functions in different cell types in homeostatic conditions. Further scRNA-seq of the fly gut will allow a number of questions to be addressed. These include changes in cell states, cell type composition and transcriptomes in the context of regeneration, various mutant backgrounds, and disease models, such as Yorkie induced intestine tumor model (Kwon et al., 2015). In addition, as female midguts are larger and longer than the male midgut and ISCs proliferation activity is higher in females, scRNA-seq will allow analysis of the basis of physiological and organ function sex differences (Hudry et al., 2016). Furthermore, during aging, changes in ISCs proliferation, regeneration capacity, innate immune and inflammatory response and tissue integrity occurs, which can be analyzed using scRNA-seq. Altogether, future scRNA-seq will provide the fundamental understanding the changes in cell states and interplay among cell types and disease.

## Acknowledgements

The authors thank Mandovi Chatterjee and Sarah Boswell from the Single Cell Core at ICCB-Longwood screening facility for InDrop encapsulation and library preparation. We thank the Transgenic RNAi Project (TRiP) and the Bloomington Drosophila Stock Center (BDSC) for providing fly stocks, the Developmental Studies Hybridoma Bank (DSHB) for monoclonal antibodies. We thank Luis Felipe Mirabella Escobedo and Henrique Camara for their help in gut dissection, Li He, Afroditi Petsakou, Stephanie Mohr, Qing Lan and members of the Perrimon laboratory for discussion and critical comments on the manuscript. This work was supported by Jane Coffin Childs Foundation (RJ Hung), TRIP R01GM084947 and DRSC R01GM067761. NP is an investigator of the Howard Hughes Medical Institute. Work by RK and SHS at the Harvard Chan Bioinformatics Core was funded by the Harvard Medical School Tools and Technology Committee and with the support of Harvard Catalyst, The Harvard Clinical and Translational Science Center (NIH award #UL1 RR 025758 and financial contributions from participating institutions). The content is solely the responsibility of the authors and does not necessarily represent the official views of the National Center for Research Resources or the National Institutes of Health. Portions of this research were conducted on the Orchestra High Performance Compute Cluster at Harvard Medical School. This NIH supported shared facility consists of thousands of processing cores and terabytes of associated storage and is partially provided through grant NCRR 1S10RR028832-01. See http://rc.hms.harvard.edu for more information.

## Methods

### Single-cell suspension preparation

*Drosophila* guts were dissociated to single cells as previously described (Dutta et al., 2015) with a few modifications. Guts were dissected from 7-day old adult *esg-sfGFP/+, pros-Gal4>RFP/+* females. After pulling out the gut, the crop and midgut/hindgut junction (where the Malpighian tubules branch out of the gut) and Malpighian tubules were removed. Five flies at a time were dissected and immediately transferred into cold PBS containing 1% BSA to avoid exposing the midgut tissue to room temperature for a long period of time. Once 40 guts were dissected, they were transferred to a dissection plate and chopped into small pieces using a razor blade. These small fragments were immediately transferred to an eppendorf tube containing 400 μl of 1 mg/ml elastase/PBS solution (Sigma E0258) and incubated on a shaker at 27 °C for 30 min. 1% of BSA in PBS (final concentration) was used to stop the digestion reaction and prevent cells from aggregating. The cell suspension was passed through 100 μm and then a 25 μm cell strainer and loaded on the top of Optiprep with density gradient 1.12 g/ml. Viable cells were isolated from the top layer of the sample after centrifugation at 800xg for 20 min. Cell viability and number were assessed by 0.4% trypan blue and hemocytometer. Cells (>160 cells/μl) were encapsulated at the Single Cell Core from ICCB-Longwood screening facility with inDrop (Zilionis et al., 2017). Reverse transcription and library preparation were done at the same facility as previously described (Zilionis et al., 2017).

### High-throughput sequencing

Before sequencing, the fragment size of each library was analyzed on a Bioanalyzer high sensitivity chip. Libraries were diluted to 1.5 nM and quantified by qPCR using primers against p5-p7 sequence. InDrop libraries were sequenced on a Nextseq 500 instrument (Illumina) with following sequencing parameters: 61 bp read 1 – 8 bp index 1 (i7) – 8 bp index 2 (i5) – 14 bp read 2.

### Datasets processing

Reads were processed using the inDrop v3 pipeline implemented in bcbio-nextgen version 1.0.5a0-9ae5245 (https://github.com/bcbio/bcbio-nextgen). Briefly, dual cellular barcodes, sample barcodes and UMIs were detected using umis (Svensson et al., 2017) correcting cellular and sample barcodes of edit distance one from expected barcodes. Cells with less than 500 total reads assigned to them were discarded. Reads were aligned to the *Drosophila melanogaster* transcriptome from FlyBase, version FB2017_03 (Gramates et al., 2017) using RapMap (Srivastava et al., 2016), assigning evidence `e` of 1/N for each read aligning to N different transcripts. Evidence was summed across all transcripts of a gene, and thresholded at a minimum evidence of 1. Quality control was performed using bcbioSingleCell (https://github.com/hbc/bcbioSingleCell), filtering out poor quality cells by keeping cells with the following metrics:

1. >= 500 UMI counts per cell
2. >= 200 genes detected per cell
3. log10(genes detected)/log10(UMI counts per cell) >= 0.65

Following these criteria, a total of 1753 cells across two replicates remained. Next, we performed the follow up analysis using Seurat (Butler et al., 2018). Prior to PCA and clustering, we regressed out the total UMI per cell and the number of genes detected per cell. As we observed little effect of the cell cycle, we did not correct for cell cycle stages. We ran PCA on the matrix of UMI corrected counts using the most variable genes and clustered the PCA dimensions with a cluster resolution parameter of 0.8, which identified 12 separate groups. We visualized the clusters using t-SNE and identified the cell types in each cluster using sets of known markers (Table S2).

### Additional marker genes selection

While Seurat efficiently identified marker genes in each cluster, they fail to identify genes expressed in a small number of cells and at low level. Thus, to recover additional markers, we added genes that met any one of the following three criteria: (1) The gene is expressed in more than 10 cells. In addition, the gene is expressed in >60% of cells in one cluster but less than 20% of cells in other clusters. (2) The gene is expressed in more than 10 cells. In addition, the gene is expressed in >80% of cells in multiple clusters and >20% of cells in all clusters. (3) The gene is expressed in a small number of cells (3-10 cells) and all of the cells that express the gene are in the same cluster.

### Gene set enrichment analysis

We performed gene set enrichment analysis on marker genes for each cluster using a program written in house. Gene sets of major functional groups were collected from the GLAD database (Hu et al., 2015), and gene sets of metabolic pathways were from the KEGG database (Kanehisa et al., 2017; Kanehisa and Goto, 2000; Kanehisa et al., 2016) while the gene sets of cellular compartment annotation were from gene ontology annotation (Ashburner et al., 2000; The Gene Ontology, 2017). In addition, the transcriptional target genes of major signaling pathways were assembled manually from the literature. P value enrichment was calculated based on the hypergeometric distribution.

### Comparison between scRNAseq and Flygut-seq data sets

To compare our scRNAseq data set with Flygut-seq (http://flygutseq.buchonlab.com/), we first identified the genes preferentially expressed in ISCs, EEs and ECs respectively in the Flygut-seq data set. Because there are no defined cell type specific genes from Flygut-seq, we generate the cell type specific gene list by calculating the ratio of expression level in each cell type vs all cell types. Specifically, the gene expression fold change cut off for ISCs and ECs is 1.6 and for EEs is 3.6. Next, we analyzed the distribution of the top 100 genes of each cell type defined by our scRNAseq data set in Flygut-seq. 89% of ISCs, 90% of EEs and 61% of ECs preferentially expressed gene were also identified by Flygut-seq.

### Cell type comparison between *Drosophila* scRNAseq and mammalian intestinal and airway epithelium

For intestine cell types, we used the cell type specific markers reported using 3p droplet (Haber et al., 2017). For some cell types eg. immature proximal enterocytes, only one marker met the criteria used in Haber et al. (fold change cutoff of 0.5 and FDR 0.05). If the number of markers was too small (less than 5), we expanded the marker gene selection using looser criteria (fold change cutoff of 0.25 instead of 0.5). We used DIOPT (release 7, score>3) (Hu et al., 2011) to map the mouse genes to *Drosophila* orthologs. We also used the mouse airway epithelium markers genes identified in (Montoro et al., 2018) with the same DIOPT criteria to map mouse genes to *Drosophila* orthologs. P value of enrichment was calculated based on the hypergeometric distribution.

### STREAM

We ran the Single-cell Trajectory Reconstruction (STREAM) for pseudotime analysis (http://stream.pinellolab.org/). We selected the default parameters, except “feature selection” for which we chose “all” to keep all genes. After input of all the expression data, the trajectories were built and displayed as a subway map plot. S2 was selected as a start state because we identified it as ISCs/EBs via our marker analysis. The genes of interest are visualized as a STREAM plot. STREAM also detected “diverging genes”, i.e. genes important in defining branching points that are differentially expressed between diverging branches and “transition genes”, i.e. genes for which their expression correlate with the cell pseudotime on a given branch.

### Fly genetics

The following strains were obtained from the Bloomington *Drosophila* Stock Center: *klu mi05554* (BL44148), *y v; attp2* (landing site only, BL36303), *y v; UAS-LucRNAi, attp2* (BL31603), *UAS-klu RNAi* (BL28731 and BL64967), *zip-Gal4* (BL48187), *lola-Gal4* (BL45325), *lrch-Gal4* (BL63768), *insc-Gal4* (BL25773), *UAS-mCD8.ChRFP* (BL27392). Strains from the Perrimon lab stock collection are: *esgGal4 UAS-GFP tubGal80^ts^ (EGT), esg-lacZ* (*esg*^*k00606*^), *Su(H)-LacZ, esg-sfGFP* (generated by David Doupe), *esg-sfGFP, UAS-mCherryCAAX* (generated by Li He), Flies were reared on standard cornmeal/agar medium in 12:12 light:dark cycles at 25°C unless noted otherwise. Fly food was changed every two days to keep fresh. Conditional expression in adult flies using *tubGal80ts* was achieved by maintaining flies at 18°C until four days after eclosion, and then shifting young adults to 29°C for 1 week. Each vial typically consisted of 10 females and 5 males were in one vial.

### HStaining and fluorescence imaging

7-day old (or unless otherwise indicated) adult female midguts were dissected in PBS, fixed with PBS containing 4% paraformaldehyde for 1hr. Sample were rinsed with PBS three times and incubated with PBST (PBS with 0.2% Triton X-100) containing 5% of normal donkey serum for 45 min. Midguts were stained at 4°C overnight with the following primary antibodies in PBST: Primary antibodies used were mouse anti-Prospero (1:100; #MR1A, Developmental Studies Hybridoma Bank), chicken anti-GFP (1:200; Aves, GFP-1020), mouse anti-β-galactosidase (1:1000; Promega). Fly midguts were washed with PBST three times and incubated with secondary antibodies (1:1000) and DAPI (1:1000) in PBST at room temperature for 1hr in the dark. Secondary antibodies were donkey anti-mouse, anti-rabbit and anti-chicken conjugated to Alexa-488, Alexa-594 and Alexa-647 (Molecular probes). Fly midguts were then washed with PBST three times, mounted in Vectashield (Vector Laboratories). Images were captured with a Zeiss LSM780 confocal microscope equipped with 20x oil lens. All images were adjusted and processed using Fiji.

### RT-qPCR

10 midguts from females were dissected, placed into TRIzol reagent (Thermo Fisher), and homogenized with Bullet Blender (Next Advance, Inc.) RNA was extracted using Direct-zol (Zymo research), converted to cDNA using SuperScript II reverse transcriptase (Invitrogen). RT-qPCR was performed using SYBR Green Supermix (Bio-Rad) with *rp49* as an internal control. Primers used for RT-qPCR are shown below:

AstA_FW: 5’-GACCTGGCCGACAGAACAAG

AstA_RV: 5’-AAAGTTGAAGGGTTGCGGAC

Tk_FW: 5’-CAATTCCTTTGTGGGGATGCG

Tk_RV: 5’-CTGCTGTTTTCCTCTCAAGTCAT

rp49_FW: 5’-ATCGGTTACGGATCGAACAA

rp49_RV: 5’-GACAATCTCCTTGCGCTTCT

